# Environmental Exposures Influence Nasal Microbiome Composition in a Longitudinal Study of Division I Collegiate Athletes

**DOI:** 10.1101/2020.02.13.946475

**Authors:** Oliver Kask, Shari Kyman, Kathryn A. Conn, Jenny Gormley, Julia Gardner, Robert A. Johns, J. Gregory Caporaso, Dierdra Bycura, Jay T. Sutliffe, Emily K. Cope

## Abstract

**Background:** The anterior nares host a complex microbial community that contributes to upper airway health. Although the bacterial composition of the nasal passages have been well characterized in healthy and diseased cohorts, the role of prolonged environmental exposures and exercise in shaping the nasal microbiome in healthy adults is poorly understood. In this study, we longitudinally sampled female collegiate Division I athletes from two teams experiencing a similar athletic season and exercise regimen but vastly different environmental exposures (Swim/Dive and Basketball). Using 16S rRNA gene sequencing, we evaluated the longitudinal dynamics of the nasal microbiome pre-, during-, and at the end of the athletic season.

**Results:** The nasal microbiota of the Swim/Dive and Basketball teams were distinct from each other at each time point sampled, driven by either low abundance (Jaccard, PERMANOVA p<0.05) or high-abundance changes in composition (Bray-Curtis, PERMANOVA p<0.05). The rate of change of microbial communities were greater in the Swim/Dive team compared to the Basketball team characterized by an increase in *Staphylococcus* in Swim/Dive and a decrease in *Corynebacterium* in both teams over time.

**Conclusions:** This is the first study that has evaluated the nasal microbiome in athletes. We obtained longitudinal nasal swabs from two gender-matched teams with similar age distributions (18-22 years old) over a 6 month period. Differences in the microbiota between teams and over time indicate that chlorine exposure, and potentially athletic training, induced changes in the nasal microbiome.

## Background

The human microbiome is defined as the collection of genomes contained within the bacteria, viruses, and fungi that inhabit nearly every niche in the body [1,2]. Recent advances in sequencing and bioinformatics technologies have vastly expanded our collective understanding of the contributions of resident microbiota to host health. Mechanistically, bacterial, viral, and fungal microbiota interact with the host to influence health or disease status through direct interaction with immune cells, indirect immune modulation through the production of metabolites such as short-chain fatty acids (SCFAs), and by contributing to mucosal epithelial barrier integrity [3–7]. The role of upper airway microbiota, including nasal microbiota, to host health is of interest, however, the dynamics of the nasal microbiota over time and under distinct environmental conditions are not well understood.

The nasal passages are a first site of contact with the external environment. In this role, the nasal microbiome has been associated with upper and lower airway health status, including upper respiratory tract infections, asthma, and *Staphylococcus aureus* pathogen carriage [8,9]. While a common feature of nasal microbiome composition is the dominance of *Corynebacterium, Staphylococcus*, and *Propionibacterium*, recent studies have identified that ecological succession and colonization patterns within individuals are associated with host health status [8–10]. A longitudinal study of 200 children demonstrated that early life succession patterns of the nasopharyngeal microbiome relate to risk of acute respiratory infection (ARI). Children with nasopharyngeal communities dominated by *Streptococcus*, *Moraxella*, or *Haemophilus* were associated with ARI, and early colonization with *Streptococcus* was a strong predictor of asthma development by 5-10 years of age [11]. The nasal passages are also an important niche for human pathobionts. Up to 20% of humans are carriers of *S. aureus* in their nares and nasal microbiota colonization patterns can influence *S. aureus* carriage [12]. *Corynebacterium accolens* and *Corynebacterium pseudodiptheritcum* presence in the nasal cavity are negatively associated with persistent *S. aureus* carriage, and these organisms compete *in vitro* [13]. *C. accolens* also exhibits anti-pneumococcal activity through the production of free fatty acids from nostril triacylglycerols [14]. Thus, understanding host and environmental factors underlying nasal microbiota structure are important for respiratory health.

Competitive athletes, especially those who are chronically exposed to allergens, pollutants, or environmental stressors, are susceptible to upper and lower airway damage leading to airway hyperresponsiveness [15]. Chlorine is an inexpensive reagent used in most swimming pools as a disinfectant and it reacts with organic matter to produce derivatives such as chloramines and gaseous nitrogen trichloride (NCl3) [16]. Although concentrations of chlorine byproducts are regulated by the World Health Organization, chronic exposure may lead to damage or irritation of the airway mucosa. Indeed, competitive, elite swimmers have a higher prevalence of upper respiratory symptoms including rhinitis and allergen sensitization [15,17,18]. A small study of 69 elite swimmers and non-swimmers demonstrated that up to 74% of swimmers report rhinitis compared to 40% of non-swimmers [17]. Upper airway symptoms can lead to poor performance and decreased quality of life [19]. Recent studies have shown that nasal microbiome composition is directly related to rhinitis and asthma [20–22]. Thus, we hypothesized that the composition or diversity of the nasal microbiota during the course of an athletic season would be altered in competitive elite swimmers compared to age- and gender-matched athletes who are not exposed to chlorine or chlorine byproducts.

Our understanding of factors that alter the nasal microbiota continue to be investigated but gaps still remain. For example, whether common environmental exposures alter microbial dynamics or composition in healthy, active adults is poorly understood. Temporal changes in the nasal microbiome in age and gender-matched groups exposed to distinct environmental pressures have not been extensively studied. Here, we evaluate temporal dynamics of the bacterial microbiome in NCAA Division I collegiate athletes chronically exposed to chlorinated water (Swim and Dive) compared to those who are unexposed to chlorine (Basketball).

## Results

### Participant Characteristics

A total of 47 subjects ages 18-22 participated in this study. All subjects were female collegiate-level NCAA Division I athletes during their competitive seasons. To determine whether prolonged chlorine exposure altered nasal microbiome composition, we recruited individuals from swim and dive and compared to female subjects on a basketball team so that microbiome comparisons matched to the same age range and athletic aptitude. Participants provided 2-5 nasal swabs at approximately one-month intervals (Fig. 1).

**Fig. 1.**
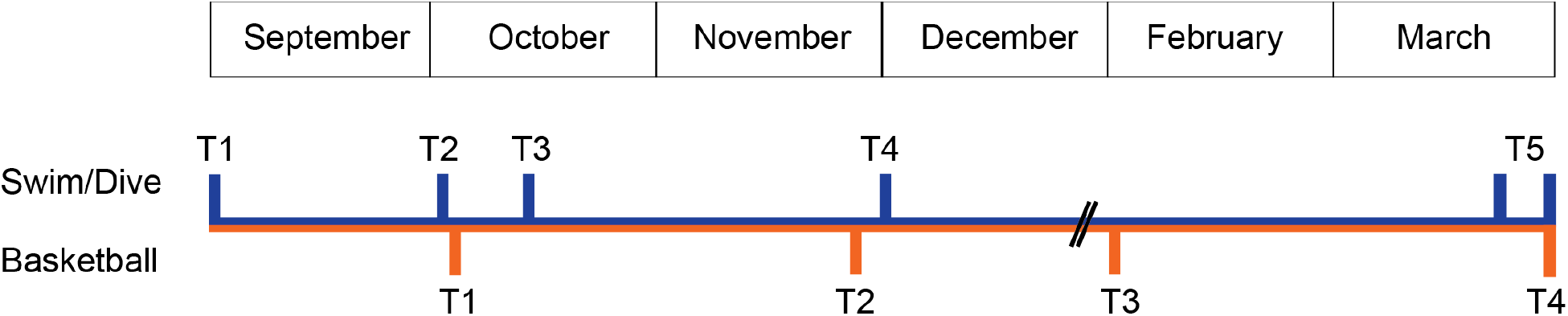
Sampling Timeline. Sampling schematic for athletes on a Swim/Dive or Basketball team. Swim/Dive participants took up to five nasal samples from September 2017 to March 2018. Basketball participants took up to four nasal swabs from October 2017 to March 2018.

Nasal swabs from participants yielded a total of 16,971,801 16S rRNA gene sequences (median per sample: 69,005, range per sample: 1,500-554,058). A total of 178 samples were sequenced, three samples were removed due to low sequence depth (<2,771 sequences/sample). Therefore, 175 specimens were included in this study.

### Compositional alterations in the nasal microbiome in chlorine exposed athletes

Alpha diversity, measured by richness (observed ASVs) did not significantly change in chlorine-exposed (Swim and Dive) vs. chlorine-unexposed (Basketball) collegiate athletes over time (Fig. 2A, Additional Figure 1). Microbial volatility analysis was used to evaluate the rate of change in nasal microbiome richness (observed ASVs) over successive time points within the same individual. There was no significant change in bacterial richness at the final time point when compared to sampling baseline time point 1 [Fig. 2A, Additional Figure 1, p=0.394 (Swim/Dive) and p=0.935 (Basketball), Wilcoxon Signed Rank Test] or to prior timepoints (Fig. 1B, Additional Figure 1, p>0.05, Wilcoxon Signed Rank Test).

**Figure 2.**
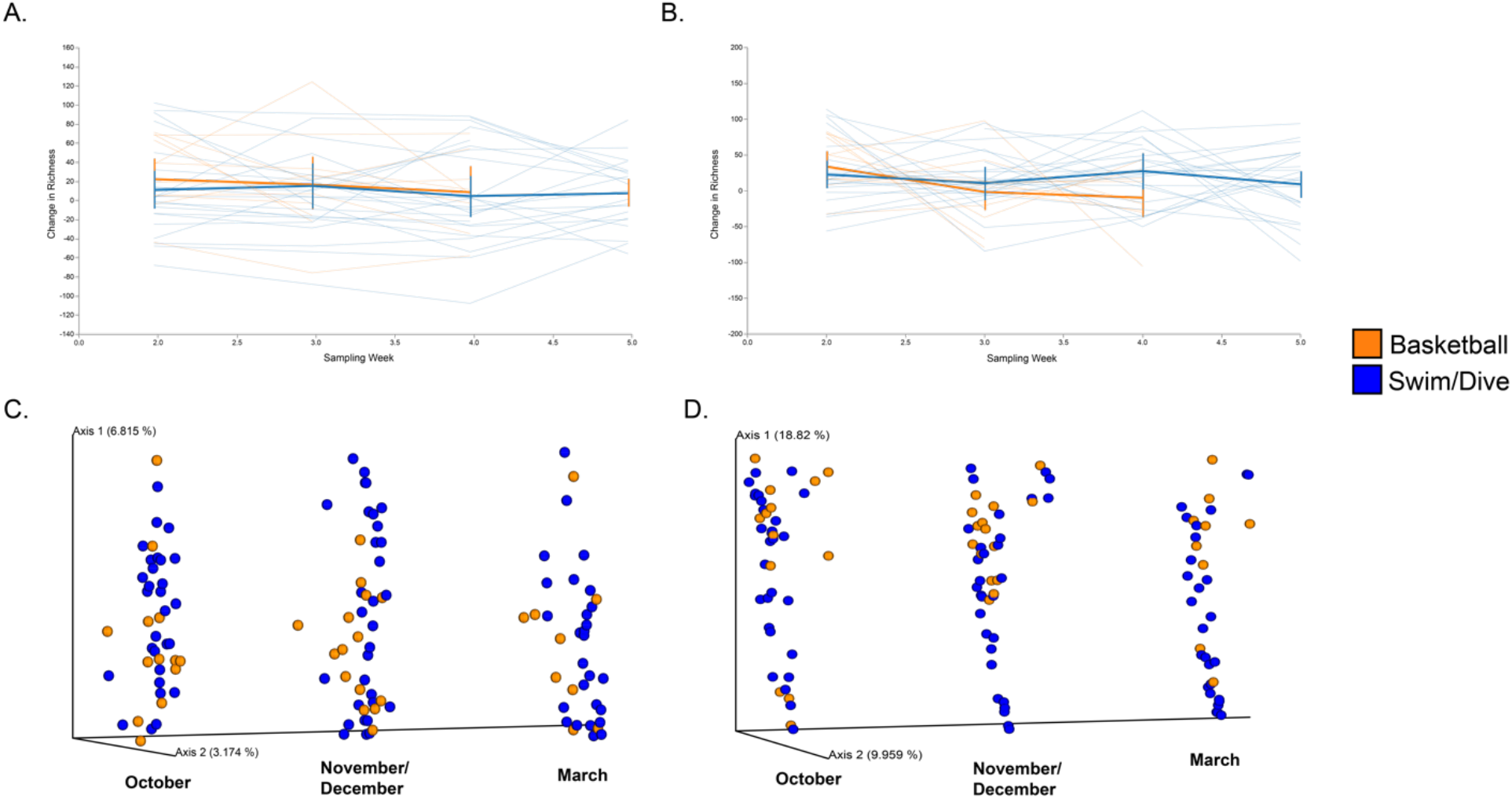
Temporal Changes in Richness and Beta-Diversity. Diversity analysis across athletic teams. Magnitude of change in bacterial richness compared to each prior sampling point [first differences, (A)]. and to baseline richness [first sampling per team; (B)]. demonstrates no significant difference across athletic team or sampling time point (p>0.05, Kruskal Wallis). Faded thin lines demonstrate the longitudinal trajectory of individuals, thick colored lines represent the mean change and standard deviation across non-chlorine exposed (Basketball) and chlorine-exposed (Swim/Dive) athletes. PCoA of Jaccard (C) and Bray-Curtis (D) distance matrices showing significant clustering by team across each sampling points matched by month (x-axis).

Multivariate analysis (PERMANOVA) of nasal bacterial beta diversity demonstrated a significant change in the athlete nasal microbiome composition between teams. To account for repeated measures, we filtered the feature table by sampling time point corresponding to overlapping season between teams [Fall (October, temperature in Flagstaff on day of sampling: 73 °F), Winter (November/December, temperatures 42 and 52 °F), and Spring (March, temperature 66 °F)]. We chose to perform this analysis at timepoints that corresponded to overlapping months, instead of sampling (see Fig. 1), to control for seasonal variation in nasal microbiome beta diversity, which has been previously described [23,24]. Indeed, when timepoints were compared without correction for season, we observed less robust clustering by team when sampling number spanned seasons (when we compare T3 for Swim/Dive in October v. T3 Basketball in January, and T4 Swim/Dive in November v. T4 Basketball in March; Additional Table 1).

Significant differences in nasal microbiome composition between individuals, grouped by athletic team, were observed using abundance-weighted (Bray-Curtis) and unweighted (Jaccard) beta diversity metrics. In Fall (October) we observed robust clustering by team with Jaccard (p=0.002, PERMANOVA, Table 2, Fig. 2C, Additional Figure 2), and Bray-Curtis (p=0.042, PERMANOVA, Table 2, Fig. 2D) metrics. In the winter (November/December), we observed significant differences when an unweighted metric was used (Jaccard, p=0.021, PERMANOVA, Table 2) but not when a weighted metric was used (Bray-Curtis, p=0.051, PERMANOVA, Table 2) indicating that, at this time point, low abundance features are joining or leaving the community (either physically, or their abundance is changing in a way that is relevant to our detection threshold). At the final overlapping time point in spring (March), bacterial communities between the two teams were significantly different when using a weighted metric (Bray-Curtis, p=0.031, PERMANOVA, Table 2), but not an unweighted metric (Jaccard, p=0.284, PERMANOVA, Table 2). Together, these results suggest that nasal microbiome alpha-diversity (richness and evenness) is relatively stable, but that compositional changes in bacterial communities are apparent between athletic teams, possibly due to chronic exposure to chlorinated water. That we observe the greatest compositional difference between teams at the beginning of the athletic season suggests that training and exercise generally may have a consistent impact on the microbial composition of the nasal cavity independent of sport.

**Table 2.** Multivariate analysis of beta diversity dissimilarity metrics by athletic team at each sampling point

| Distance Metric | Sampling Time Point | pseudo-F | PERMANOVA<br>p value |
| --- | --- | --- | --- |
| **Bray-Curtis** | **October** | **1.82** | **0.042** |
| **Jaccard** | **October** | **1.32** | **0.002** |
| Bray-Curtis | November/December | 1.69 | 0.051 |
| **Jaccard** | **November/December** | **1.35** | **0.021** |
| **Bray-Curtis** | **March** | **2.18** | **0.031** |
| Jaccard | March | 1.05 | 0.284 |

### Environmental exposures are associated with reduced bacterial microbiome stability

Next, we sought to determine how microbiome composition changes within an individual over time and whether chronic exposure to chlorinated water alters microbial stability, defined here as the magnitude of change in community dissimilarity metrics within an individual. We used q2-longitudinal[25]. to examine how community dissimilarity metrics changed in each individual between successive time points in each athletic team (Fig. 3). The magnitude of change in Jaccard and Bray-Curtis dissimilarity indices increased at the final timepoint (March) in the Swim/Dive team, indicating that the rate of change of beta diversity was greater in individuals exposed to chlorinated water (Fig. 3A, Fig. 3B). Bray-Curtis and Jaccard indices remained stable (Fig. 3A) or decreased (Fig. 3B) in the basketball team, indicating more similar bacterial communities within an individual over time. The magnitude of change in Jaccard dissimilarity metric was significantly greater in individuals on the Swim/Dive team than in individuals on the Basketball team between the first and final time points (Jaccard p=0.008, Mann-Whitney U; Bray-Curtis p=0.466, Mann-Whitney U).

**Figure 3.**
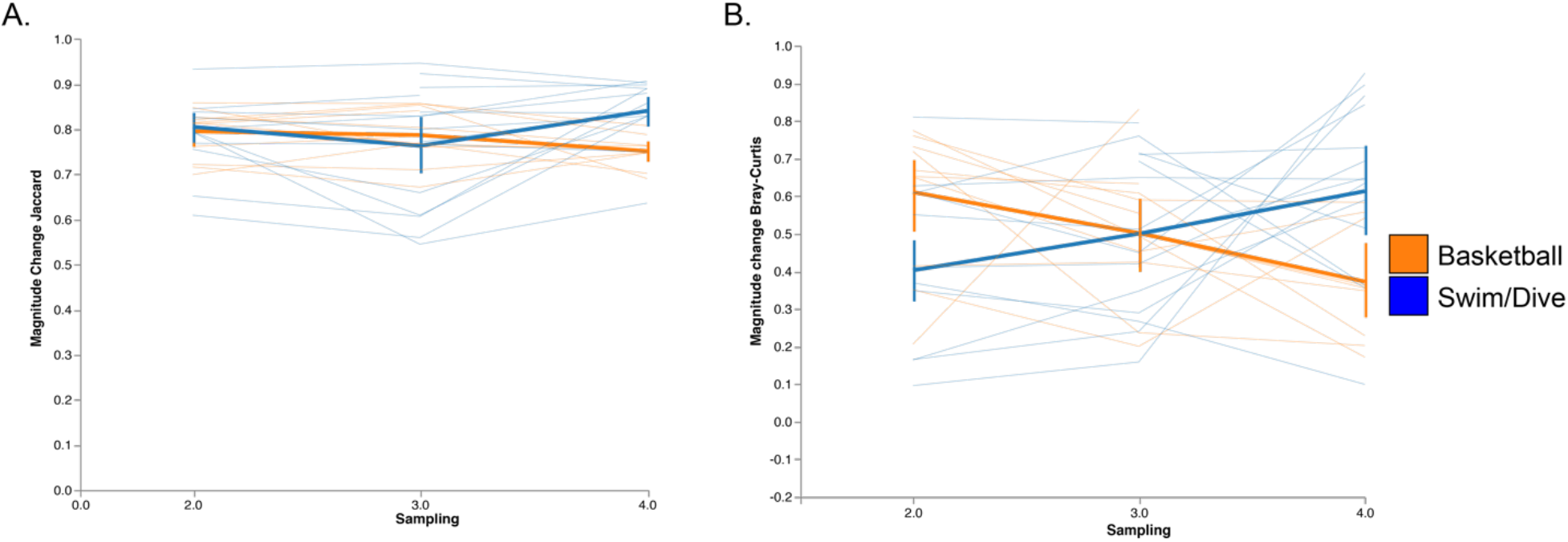
Magnitude Change in Jaccard and Bray-Curtis Over Successive Timepoints. Longitudinal change in Jaccard (A) and Bray-Curtis (B) distances between successive samples (first distances) from chlorine-exposed (Swim/Dive) and unexposed (Basketball) athletes. Athletes who were not exposed to chlorine during the athletic season had more similar nasal microbiota when measured using Jaccard but not Bray-Curtis dissimilarity metric, at the end of the sampling period compared to the baseline sample whereas athletes who were chronically exposed to chlorine had less similar nasal microbiota compared to baseline and prior sampling timepoints (Jaccard, p=0.02, Kruskal Wallis; Bray-Curtis p=0.45 Kruskal Wallis).

To identify important features (e.g. taxa) that change in abundance over time in each athletic team, we used a supervised learning regressor as implemented in q2-longitudinal feature volatility analysis (Random Forest Regressor, model accuracy p=0.000003, Mean Squared Error=0.300, r^2^=0.622 [25]). Two low-abundance features were strong predictors of sampling time point in each team: TM6 increased over time (feature importance score=0.180, net average change=0.0002) and unclassified *Bacillaceae* decreased over time (feature importance score=0.079, net average change=-0.0922). Two of the highest abundance features were *Staphylococcus* and *Corynebacterium* (Additional Figure 3) and were also strong predictors of sampling time point. *Staphylococcus* was ranked third in feature importance and on average, increased over time with a higher rate of increase in individuals on the Swim/Dive team (feature importance score=0.064, net average change=0.1442; Fig. 4A). The second most abundant feature, *Corynebacterium*, decreased in each athletic team over time in both teams (feature importance score=0.008, net average change=-0.1070; Fig. 4B). These results support our prior finding that compositional differences are driven by high-abundance taxonomic changes between athletic teams at the final sampling time point.

**Figure 4.**
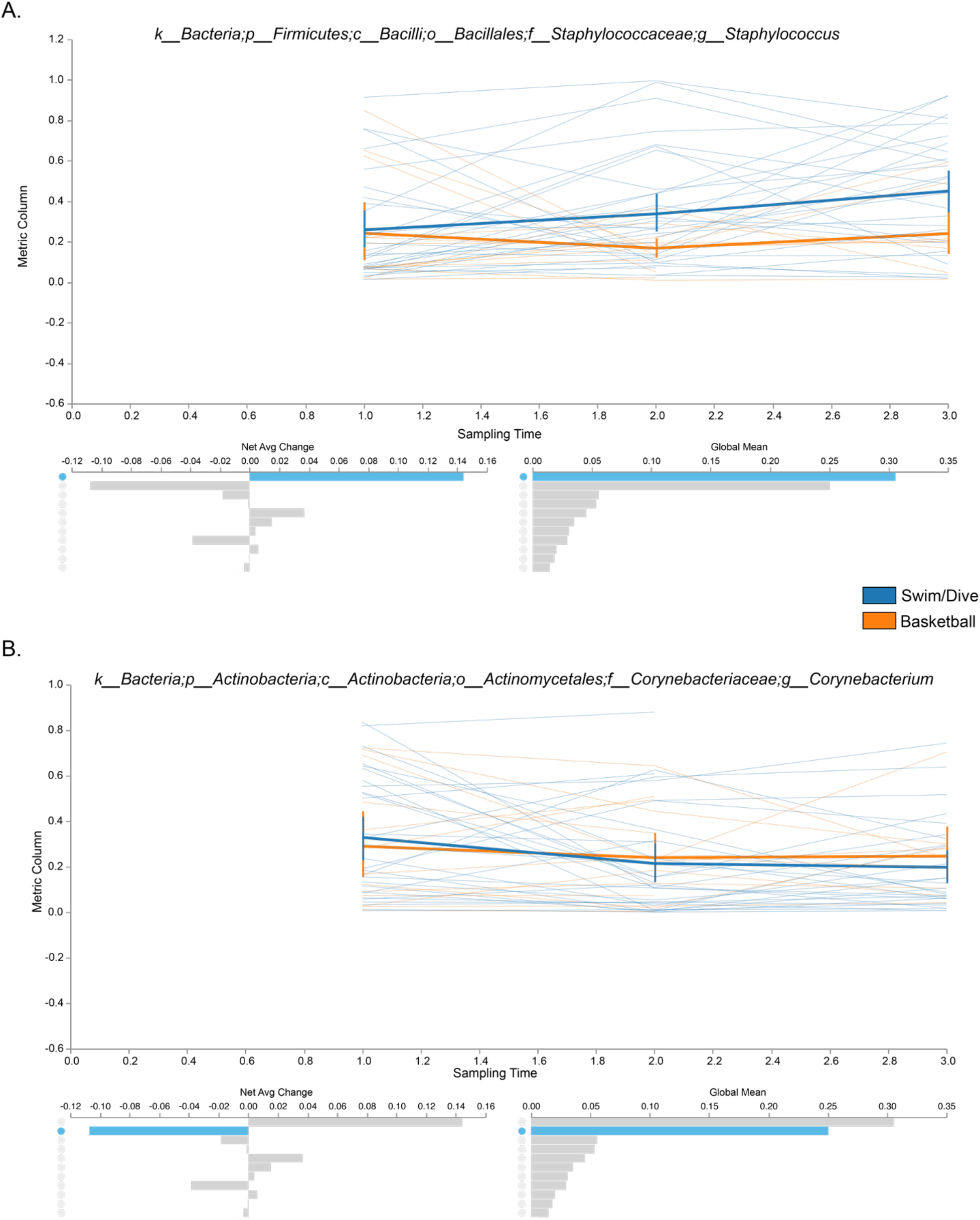
Random Forest Regression Identified Change of Important Features Over Time. Random Forest regression of feature abundance over time. Two high abundance features were predictive of sampling time point. A) *Staphylococcus* abundance increased over time in participants on the Swim/Dive team, but remained relatively stable in participants on the Basketball team (feature importance score=0.064, net average change=0.1442). B) *Corynebacterium* abundance decreased over time in participants on both teams (feature importance score=0.008, net average change= −0.1070).

Evaluating changes in individual features is important, however microbial communities exist within a complex ecological framework comprised of competing, commensal, and mutualistic interactions. Thus, microbes do not change in response to environmental pressures in isolation. To determine whether networks of bacteria change over time in athletes exposed to chlorinated water, we used a nonparametric microbial interdependence test (NMIT) [26]. as implemented in q2-longitudinal [25]. to determine whether distinct networks of interdependent ASVs change within each group (swim and dive v. basketball). To ensure robust microbial interdependence networks, we excluded individuals with fewer than 3 sampling timepoints. NMIT demonstrated the microbiome of swimmers and divers exhibited similar temporal characteristics, or similar networks of bacteria changing, when compared to individuals on a basketball team (Additional Figure 4, PERMANOVA pseudo-F 1.22, p=0.022). These results suggest that networks of microbiota may change in response to chronic chlorine exposure.

Athletes on the Swim/Dive and Basketball teams were asked to complete a Sinonasal Outcomes Test (SNOT-22), a validated questionnaire that surveys nasal, behavioural, and sleep symptoms. We evaluated microbiome alpha diversity and nasal symptoms (the sum of SNOT-22 questions 1-7). Higher values indicate worse nasal symptoms. There was not a significant difference in nasal symptom scores between teams (p=0.117, Welch’s t-test of averaged SNOT-22 score for each individual) though a larger sample size may be needed to identify this association. We also evaluated whether an increase in nasal symptoms was correlated with decreased alpha diversity using repeated measures correlation (rmcorr) in R [27]., as has been demonstrated in the sinonasal cavity in sinusitis [28]. We observed a weak but non-significant negative correlation overall (r^2^= −0.117, p=0.261) and in the Swim/Dive team (r^2^=0.029, p=0.868, Additional Figure 5). We observed no correlation between SNOT-22 and richness in individuals on the basketball team (r^2^= 0.029, p=0.868). Our results therefore do not illustrate a relationship between SNOT-22 scores and alpha diversity, but we expect that a larger sample size including individuals with diagnosed upper respiratory disease might elucidate a relationship.

## Discussion

We demonstrated that healthy, Division I NCAA athletes who are chronically exposed to chlorinated water (Swim and Dive) have distinct microbial community composition, but not alpha diversity, during the active season when compared to athletes who are not exposed to chlorinated water (Basketball). Both teams were comprised of healthy, female athletes, aged 18-22. The nasal microbiota of individuals on the Swim and Dive team were compositionally less similar to their baseline sample over time, while the nasal microbiota of individuals on the Basketball team was compositionally more similar to their baseline sample over time. This was expected, since the Swim and Dive nasal microbial community was exposed to chronic antimicrobial pressures via chlorination and chlorine byproducts, presumably strong drivers of bacterial composition. In this study, causality of chlorine exposure alterations to the nasal microbiome could not be determined since the nasal microbiome composition of each team was distinct at the start of the study, possibly due to years of chlorine exposure prior to joining an NCAA Division I swim and dive team.

Previous studies have demonstrated an increased prevalence of allergic rhinitis, asthma, and allergies in elite athletes exposed to irritants and chlorinated water [18,29,30]. These studies are intriguing, but are limited in scope and unable to detect specific mechanistic pathways by which chronic exposure to low levels of chlorine and chlorine derivatives drive airway disease. It’s unclear whether athletes choose an athletic career in the pool because of underlying airway issues, such as asthma, or whether exposure to chlorine and byproducts precedes disease. We hypothesized that altered nasal microbiome composition may contribute to airway disease, as has been previously described in patients with AR and asthma [9,20,21]. While we have not mechanistically demonstrated this in the current study, our results suggest that elite athletes exposed to chlorinated water have altered microbial composition. Specifically, we demonstrated that the rate of change in Jaccard and Bray-Curtis distances increased in the Swim/Dive team over time, indicating that the nasal microbiota of participants on the Swim and Dive team became less similar than their previous sampling point as the season progressed. In contrast, the rate of change in distances was smaller in participants on the basketball team, indicating that the nasal microbiota did not change or became more similar to their previous sampling time points. When we computed differences from baseline and final sampling points, we observed significant differences in proportions of OTUs shared between samples (Jaccard); Swim/Dive athletes nasal microbiota shared fewer taxa with their baseline sample at the final time point whereas Basketball athletes nasal microbiota became shared more taxa with their baseline sample at the final time point.

We observed bacterial taxa that were predictive of sampling time point in each team. Most notably, *Staphylococcus* abundance increased in abundance in Swim/Dive participants over time, but *Corynebacterium* decreased over time in both teams, perhaps as a general consequence of athletic training. *Corynebacterium* species are commensal and potentially beneficial in the anterior nares so we were surprised to observe a decrease in *Corynebacterium* relative abundance throughout the athletic season. An antagonistic relationship between *Staphylococcus* and *Corynebacterium* has been observed in nasal microbiome and *in vitro* studies [13,28,31–33]. *Corynebacterium* can inhibit *Staphylococcus aureus* growth through secretion of anti-Staphylococcal factors [33]. and can shift the behavior of *S. aureus* toward commensalism when in a polymicrobial community by inhibiting the quorum sensing signal, accessor gene regulator (*agr*) [34]. Future studies should be aimed at understanding the health implications of increased *Staphylococcus* burden in collegiate and professional swimmers. In addition, we did not assess the effect of chlorinated, saltwater, or freshwater on the nasal microbiota, which should also be the focus of future studies.

This is the first study that has evaluated the nasal microbiome in athletes. We obtained longitudinal nasal swabs from two gender-matched teams with similar age distributions (18-22 years old) over an approximately 6 month period. The limitations of our study include uneven periods between sampling, though we were able to sample in three seasons (Fall, Winter, Spring) for each team. We also recognize that this study was performed with a relatively small sample size (n=47 total participants), so we may not have statistical power to detect true changes in the microbiome at each timepoint. Finally, we were unable to include a healthy, non-athlete cohort in this initial study. We are currently recruiting healthy, non-athletes to determine whether we observe changes in the nasal microbiome due to athletic status. Although this question has not been determined in the nasal microbiota, a recent study evaluating the skin microbiome showed major shifts in athletes playing a contact sport (roller derby) [35]. In future studies, we will also measure nasal fungal microbiota to determine whether networks of bacteria and fungi are influenced by environmental exposures. The inclusion of a healthy cohort of individuals exposed to chlorinated water represents an exciting opportunity to evaluate the stability of the nasal microbiota under environmental pressures.

## Materials and Methods

### Subject Recruitment

NCAA Division I female athletes ages 18-22 were recruited from a collegiate Swim and Dive and Basketball team in Arizona under approved IRBs 982568-4 and 982568-14. Inclusion criteria included active membership of each athletic team and general good health. No exclusionary criteria were developed for this study, as all NCAA athletes are required to complete yearly athletic physicals to screen for possible health risks. Individuals were not excluded based on prior diagnosis of airway disease, but prior physician’s diagnosis and current nasal symptoms were recorded as metadata and used in subsequent analyses. Fifteen members of the basketball team and 32 members of Swim and Dive participated. Up to 5 timepoints (a minimum of 2 timepoints) were collected per individual throughout the course of the athletic season. Nasal specimens were obtained at approximately 1-month intervals. A total of 178 nasal specimens were processed for bacterial microbiome sequencing. The active seasons for each team were similar; Swim and Dive meets occur September to March and Basketball games occur November to March. Athletes on the Swim/Dive and Basketball teams were asked to complete a Sinonasal Outcomes Test (SNOT-22), a validated questionnaire that surveys nasal, behavioral, and sleep symptoms [36].

### Nasal Specimen Collection and DNA extraction

Nasal bacterial samples were collected using a BBL CultureSwab (Becton, Dickinson and Company, Sparks, MD). Participants were instructed to swab their anterior nares with both swab tips for 10-15 seconds per nostril. Swabs were transported to a −80 °C freezer and stored until DNA extraction. Nasal samples were randomized into five extraction sets. Total DNA was extracted from nasal swabs using DNeasy PowerSoil Kit (Qiagen Hilden, Germany) using manufacturer’s protocol with one modification; to facilitate microbial lysis, swabs were incubated in lysis buffer for 10 minutes at 65 °C before sample vortexing. Resulting DNA samples were quantified on the Nanodrop 8000 Spectrophotometer (ThermoFisher Waltham, MA).

### 16S rRNA gene sequencing

Amplicon sequencing of the 16S rRNA gene for sequencing on the MiSeq Illumina platform was done using the protocol from the Earth Microbiome Project [37]. Barcoded 806R reverse primers and forward primer 515F were used to amplify the V4 region of the 16S rRNA gene [38]. Library preparation was done at the Pathogen and Microbiome Institute and sequencing was performed at the Translational Genomics Research Institute (TGen) Pathogen and Microbiome Division. PCR conditions were as follows: 2 minutes at 98 °C for 1 cycle; 20 seconds at 98 °C, 30 seconds at 50 °C, and 45 seconds at 72 °C for 30 cycles; and 10 minutes at 72 °C for 1 cycle. PCR product was purified using AMPure XP for PCR Purification (Beckman Coulter Indianapolis, IN), quantified using Qubit dsDNA HS Assay Kit (ThermoFisher Waltham, MA), and pooled at 25 ng/sample for sequencing. If extraneous DNA bands (human or mitochondrial) were present, the samples were run on an EGel size-select gel (ThermoFisher Invitrogen, Waltham, MA) prior to pooling. Extraction blank negative controls were included in each extraction set (five total) and sequenced with the pool of nasal samples. A negative control for each barcoded primer was also run and visualized on a gel. If contamination was observed in the negative well, the sample was run with a new barcoded primer.

### Microbiome Bioinformatics

16S rRNA gene sequences were analyzed using Quantitative Insights Into Microbial Ecology 2 (QIIME 2) 2019.10 [39]. Paired end sequences were demultiplexed, then denoised, chimera checked and grouped into Amplicon Sequence Variants (ASVs) based on 100% sequence identity using dada2 [40]. Sequences were aligned using MAFFT [41] and a phylogenetic tree was built using FastTree 2 [42]. Taxonomy was assigned to each ASV using a Naive Bayes classifier[43]. trained on Greengenes 13_8 99% OTUs [44]. Richness (observed OTUs), Faith’s Phylogenetic Diversity [45], and Shannon diversity were used to assess alpha diversity. Jaccard, Bray-Curtis, Weighted and Unweighted UniFrac [46] metrics were used to assess beta diversity. Longitudinal analysis was performed using the q2-longitudinal plugin [25]. Volatility analysis was used to determine how metrics (e.g. alpha diversity or beta diversity) changed over the sampling period [25].

### Statistical Analysis

In order to include the maximum number of longitudinal sampling points, we rarefied samples to an even sampling depth of 2,771 sequences/sample. Three samples were removed at this step due to low sequence depth. Kruskal-Wallis was used to evaluate changes in alpha diversity (richness, shannon, Faith’s PD) across each team within one time point [47]. Permutational analysis of variance (PERMANOVA) was used to compare beta diversity (Bray-Curtis, Jaccard, Weighted Unifrac, Unweighted UniFrac) across teams at each time point [48]. Pairwise comparisons in alpha diversity were made for each pair of samples between successive timepoints using a Wilcoxon Signed Rank Test as implemented in q2-longitudinal [25]. A Mann-Whitney U test was used to assess whether distance (using a beta diversity distance matrix) between successive timepoints were significantly different between each team over time. Finally, Repeated Measures Correlation using rmcorr in R version 1.2.1335 was used to assess the relationship between alpha diversity and the sum of the questions 1-7 on the SNOT-22 questionnaire, which evaluates nasal symptoms [27]. To identify important features (e.g. taxa) that change in abundance over time in each athletic team, we used a random forest supervised learning regressor with 5-fold cross validation and 100 estimator trees, as implemented in q2-longitudinal feature volatility analysis on a feature table collapsed to the genus level [25].

## Declarations

### Ethics approval and consent to participate

NCAA Division I female athletes ages 18-22 were recruited from a collegiate Swim and Dive and Basketball team in Arizona under approved IRBs 982568-4 and 982568-14.

### Consent for publication

Not applicable

### Availability of data and material

Sequence data for this study have been deposited in the NCBI Short Read Archive (SRA) under BioProject ID PRJNA605856.

## Acknowledgements

We would like to thank the athletes who provided nasal samples for this study and their coaches for their support of this project.

Authors’ information (optional)

## Author’s Contributions

OK and SK performed library preparation. OK and EKC analyzed sequence data. EKC and JGC interpreted sequence data. EKC, OK, and KAC wrote and edited the manuscript. EKC, DB, JTS and RAJ conceived and designed the study. All authors read and approved the final manuscript

## References

1. Human Microbiome Project Consortium. Structure, function and diversity of the healthy human microbiome. Nature. 2012;486:207–14.

2. Sender R, Fuchs S, Milo R. Revised Estimates for the Number of Human and Bacteria Cells in the Body. PLoS Biol. 2016;14:e1002533.

3. Li M, Wang B, Zhang M, Rantalainen M, Wang S, Zhou H, et al. Symbiotic gut microbes modulate human metabolic phenotypes [Internet].. Proceedings of the National Academy of Sciences. 2008. p. 2117–22. Available from: http://dx.doi.org/10.1073/pnas.0712038105

4. Ivanov II, Honda K. Intestinal commensal microbes as immune modulators. Cell Host Microbe. 2012;12:496–508.

5. Narushima S, Sugiura Y, Oshima K, Atarashi K, Hattori M, Suematsu M, et al. Characterization of the 17 strains of regulatory T cell-inducing human-derived Clostridia. Gut Microbes. 2014;5:333–9.

6. Smith PM, Howitt MR, Panikov N, Michaud M, Gallini CA, Bohlooly-Y M, et al. The microbial metabolites, short-chain fatty acids, regulate colonic Treg cell homeostasis. Science. 2013;341:569–73.

7. Hsiao EY, McBride SW, Hsien S, Sharon G, Hyde ER, McCue T, et al. Microbiota modulate behavioral and physiological abnormalities associated with neurodevelopmental disorders. Cell. 2013;155:1451–63.

8. Liu CM, Price LB, Hungate BA, Abraham AG, Larsen LA, Christensen K, et al. Staphylococcus aureus and the ecology of the nasal microbiome. Sci Adv. 2015;1:e1400216.

9. Pérez-Losada M, Castro-Nallar E, Bendall ML, Freishtat RJ, Crandall KA. Dual Transcriptomic Profiling of Host and Microbiota during Health and Disease in Pediatric Asthma. PLoS One. 2015;10:e0131819.

10. Wilson MT, Hamilos DL. The nasal and sinus microbiome in health and disease. Curr Allergy Asthma Rep. 2014;14:485.

11. Teo SM, Mok D, Pham K, Kusel M, Serralha M, Troy N, et al. The infant nasopharyngeal microbiome impacts severity of lower respiratory infection and risk of asthma development. Cell Host Microbe. 2015;17:704–15.

12. van Belkum A, Verkaik NJ, de Vogel CP, Boelens HA, Verveer J, Nouwen JL, et al. Reclassification of Staphylococcus aureus nasal carriage types. J Infect Dis. 2009;199:1820–6.

13. Yan M, Pamp SJ, Fukuyama J, Hwang PH, Cho D-Y, Holmes S, et al. Nasal microenvironments and interspecific interactions influence nasal microbiota complexity and S. aureus carriage. Cell Host Microbe. 2013;14:631–40.

14. Bomar L, Brugger SD, Yost BH, Davies SS, Lemon KP. Corynebacterium accolens Releases Antipneumococcal Free Fatty Acids from Human Nostril and Skin Surface Triacylglycerols. MBio. 2016;7:e01725–15.

15. Zwick H, Popp W, Budik G, Wanke T, Rauscher H. Increased sensitization to aeroallergens in competitive swimmers. Lung. 1990;168:111–5.

16. Organization WH, Others. Guidelines for safe recreational water environments. Volume 2: Swimming pools and similar environments. World Health Organization; 2006.

17. Bougault V, Turmel J, Boulet LP. Effect of intense swimming training on rhinitis in high-level competitive swimmers [Internet].. Clinical & Experimental Allergy. 2010. p. 1238–46. Available from: http://dx.doi.org/10.1111/j.1365-2222.2010.03551.x

18. Gelardi M, Ventura MT, Fiorella R, Fiorella ML, Russo C, Candreva T, et al. Allergic and non-allergic rhinitis in swimmers: clinical and cytological aspects. Br J Sports Med. 2012;46:54–8.

19. Meltzer EO. Quality of life in adults and children with allergic rhinitis. J Allergy Clin Immunol. 2001;108:S45–53.

20. Choi CH, Poroyko V, Watanabe S, Jiang D, Lane J, deTineo M, et al. Seasonal allergic rhinitis affects sinonasal microbiota. Am J Rhinol Allergy. journals.sagepub.com; 2014;28:281–6.

21. Durack J, Huang YJ, Nariya S, Christian LS, Ansel KM, Beigelman A, et al. Bacterial biogeography of adult airways in atopic asthma. Microbiome. 2018;6:104.

22. Fazlollahi M, Lee TD, Andrade J, Oguntuyo K, Chun Y, Grishina G, et al. The nasal microbiome in asthma. J Allergy Clin Immunol. 2018;142:834–43.e2.

23. Pérez-Losada M, Alamri L, Crandall KA, Freishtat RJ. Nasopharyngeal Microbiome Diversity Changes over Time in Children with Asthma [Internet].. PLOS ONE. 2017. p. e0170543. Available from: http://dx.doi.org/10.1371/journal.pone.0170543

24. Santee CA, Nagalingam NA, Faruqi AA, DeMuri GP, Gern JE, Wald ER, et al. Nasopharyngeal microbiota composition of children is related to the frequency of upper respiratory infection and acute sinusitis [Internet].. Microbiome. 2016. Available from: http://dx.doi.org/10.1186/s40168-016-0179-9

25. Bokulich N, Zhang Y, Dillon M, Rideout JR, Bolyen E, Li H, et al. q2-longitudinal: a QIIME 2 plugin for longitudinal and paired-sample analyses of microbiome data [Internet].. bioRxiv. 2017 [cited 2017 Dec 4].. p. 223974. Available from: https://www.biorxiv.org/content/early/2017/11/22/223974.abstract

26. Zhang Y, Han SW, Cox LM, Li H. A multivariate distance-based analytic framework for microbial interdependence association test in longitudinal study. Genet Epidemiol. Wiley Online Library; 2017;41:769–78.

27. Bakdash JZ, Marusich LR. Repeated Measures Correlation. Front Psychol. 2017;8:456.

28. Cope EK, Goldberg AN, Pletcher SD, Lynch SV. Compositionally and functionally distinct sinus microbiota in chronic rhinosinusitis patients have immunological and clinically divergent consequences. Microbiome. 2017;5:53.

29. Thickett KM, McCoach JS, Gerber JM, Sadhra S, Burge PS. Occupational asthma caused by chloramines in indoor swimming-pool air. Eur Respir J. 2002;19:827–32.

30. Helenius I, Haahtela T. Allergy and asthma in elite summer sport athletes. J Allergy Clin Immunol. 2000;106:444–52.

31. Laux C, Peschel A, Krismer B. Staphylococcus aureus Colonization of the Human Nose and Interaction with Other Microbiome Members. Microbiol Spectr [Internet].. 2019;7. Available from: http://dx.doi.org/10.1128/microbiolspec.GPP3-0029-2018

32. Proctor DM, Relman DA. The Landscape Ecology and Microbiota of the Human Nose, Mouth, and Throat. Cell Host Microbe. 2017;21:421–32.

33. Hardy BL, Dickey SW, Plaut RD, Riggins DP, Stibitz S, Otto M, et al. Corynebacterium pseudodiphtheriticum Exploits Staphylococcus aureus Virulence Components in a Novel Polymicrobial Defense Strategy. MBio [Internet].. 2019;10. Available from: http://dx.doi.org/10.1128/mBio.02491-18

34. Ramsey MM, Freire MO, Gabrilska RA, Rumbaugh KP, Lemon KP. Staphylococcus aureus Shifts toward Commensalism in Response to Corynebacterium Species. Front Microbiol. 2016;7:1230.

35. Meadow JF, Bateman AC, Herkert KM, O’Connor TK, Green JL. Significant changes in the skin microbiome mediated by the sport of roller derby. PeerJ. 2013;1:e53.

36. Kennedy JL, Hubbard MA, Huyett P, Patrie JT, Borish L, Payne SC. Sino-nasal outcome test (SNOT-22): a predictor of postsurgical improvement in patients with chronic sinusitis. Ann Allergy Asthma Immunol. 2013;111:246–51.e2.

37. Thompson LR, The Earth Microbiome Project Consortium, Sanders JG, McDonald D, Amir A, Ladau J, et al. A communal catalogue reveals Earth’s multiscale microbial diversity [Internet].. Nature. 2017. p. 457–63. Available from: http://dx.doi.org/10.1038/nature24621

38. Caporaso JG, Lauber CL, Walters WA, Berg-Lyons D, Huntley J, Fierer N, et al. Ultra-high-throughput microbial community analysis on the Illumina HiSeq and MiSeq platforms. ISME J. 2012;6:1621–4.

39. Bolyen E, Rideout JR, Dillon MR, Bokulich NA, Abnet CC, Al-Ghalith GA, et al. Reproducible, interactive, scalable and extensible microbiome data science using QIIME 2. Nat Biotechnol [Internet].. 2019; Available from: http://dx.doi.org/10.1038/s41587-019-0209-9

40. Callahan BJ, McMurdie PJ, Rosen MJ, Han AW, Johnson AJA, Holmes SP. DADA2: high-resolution sample inference from Illumina amplicon data. Nat Methods. Nature Publishing Group; 2016;13:581.

41. Katoh K, Standley DM. MAFFT multiple sequence alignment software version 7: improvements in performance and usability. Mol Biol Evol. 2013;30:772–80.

42. Price MN, Dehal PS, Arkin AP. FastTree 2--approximately maximum-likelihood trees for large alignments. PLoS One. 2010;5:e9490.

43. Pedregosa F, Varoquaux G, Gramfort A, Michel V, Thirion B, Grisel O, et al. Scikit-learn: Machine learning in Python. J Mach Learn Res. 2011;12:2825–30.

44. McDonald D, Price MN, Goodrich J, Nawrocki EP, DeSantis TZ, Probst A, et al. An improved Greengenes taxonomy with explicit ranks for ecological and evolutionary analyses of bacteria and archaea. ISME J. 2012;6:610–8.

45. Faith DP, Minchin PR, Belbin L. Compositional dissimilarity as a robust measure of ecological distance. Vegetatio. 1987;69:57–68.

46. Lozupone C, Knight R. UniFrac: a new phylogenetic method for comparing microbial communities. Appl Environ Microbiol. 2005;71:8228–35.

47. Kruskal WH, Wallis WA. Use of Ranks in One-Criterion Variance Analysis. J Am Stat Assoc. Taylor & Francis; 1952;47:583–621.

48. Anderson MJ. A new method for non-parametric multivariate analysis of variance. Austral Ecol. Wiley Online Library; 2001;26:32–46.

49. Lévesque B, Duchesne J-F, Gingras S, Lavoie R, Prud’Homme D, Bernard E, et al. The determinants of prevalence of health complaints among young competitive swimmers. Int Arch Occup Environ Health. 2006;80:32–9.

50. Wagner Mackenzie B, Waite DW, Hoggard M, Douglas RG, Taylor MW, Biswas K. Bacterial community collapse: a meta-analysis of the sinonasal microbiota in chronic rhinosinusitis. Environ Microbiol. 2017;19:381–92.

51. McCauley K, Durack J, Valladares R, Fadrosh DW, Lin DL, Calatroni A, et al. Distinct nasal airway bacterial microbiotas differentially relate to exacerbation in pediatric patients with asthma. J Allergy Clin Immunol [Internet].. 2019; Available from: http://dx.doi.org/10.1016/j.jaci.2019.05.035

