## Supplemental Figures for "Environmental Exposures Influence Nasal Microbiome Composition in a Longitudinal Study of Division I Collegiate Athletes"

**Additional Figures.**

**Additional Table 1.** Multivariate analysis of beta diversity dissimilarity metrics by athletic team at sampling points spanning seasons

| Distance Metric | Sampling Time Point | pseudo-F | PERMANOVA p value |
|---|---|---|---|
| Bray-Curtis | T3 (October v. January) | 1.516 | 0.089 |
| ***Jaccard*** | ***T3*** (October v. January) | ***1.171*** | ***0.047*** |
| Bray-Curtis | T4 (November v. March) | 1.084 | 0.343 |
| Jaccard | T4 (November v. March) | 1.065 | 0.226 |

**Additional Figure 1.**

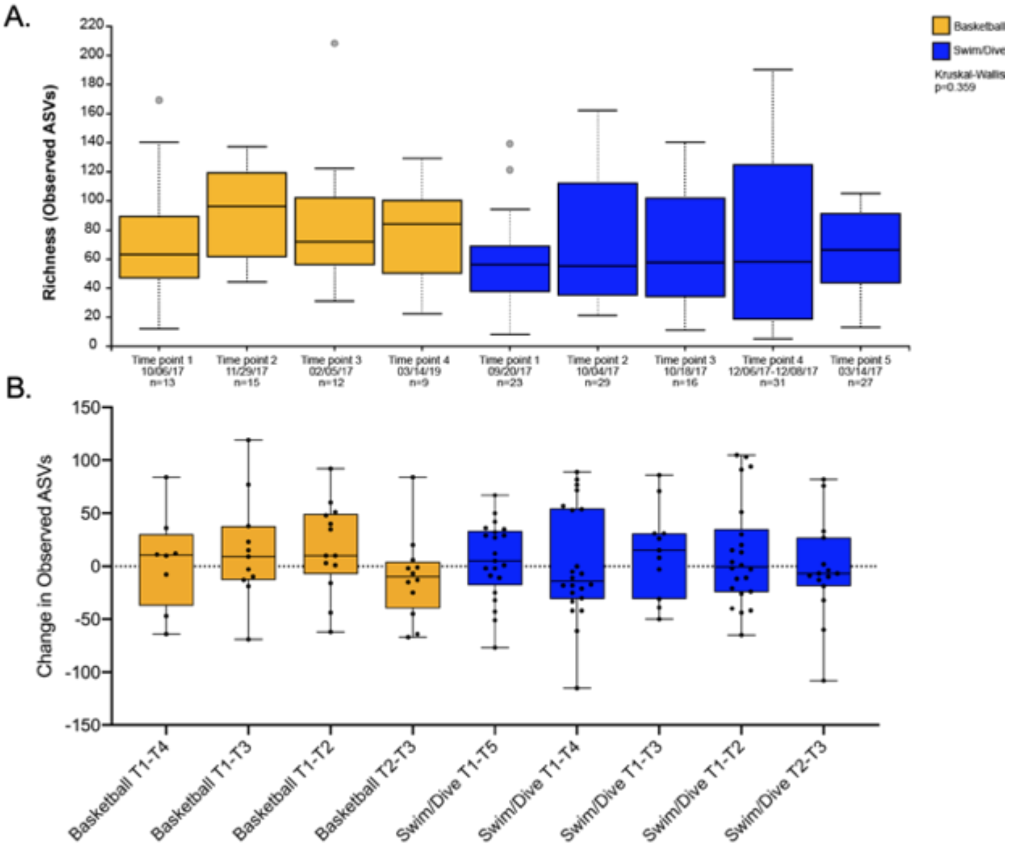


| Team | Timepoints Compared | W | p-value |
|---|---|---|---|
| Basketball | T1-T2 | 18 | 0.109 |
| Basketball | T1-T3 | 20 | 0.328 |
| Basketball | T1-T4 | 13 | 0.935 |
| Basketball | T2-T3 | 21 | 0.315 |
| Basketball | T3-T4 | 12 | 0.233 |
| Swim/Dive | T1-T2 | 105 | 0.485 |
| Swim/Dive | T1-T3 | 22 | 0.328 |
| Swim/Dive | T1-T4 | 124 | 0.935 |
| Swim/Dive | T1-T5 | 91 | 0.394 |
| Swim/Dive | T2-T3 | 52 | 0.649 |
| Swim/Dive | T3-T4 | 39 | 0.233 |
| Swim/Dive | T4-T5 | 172.5 | 0.939 |

**Additional Figure 2.**

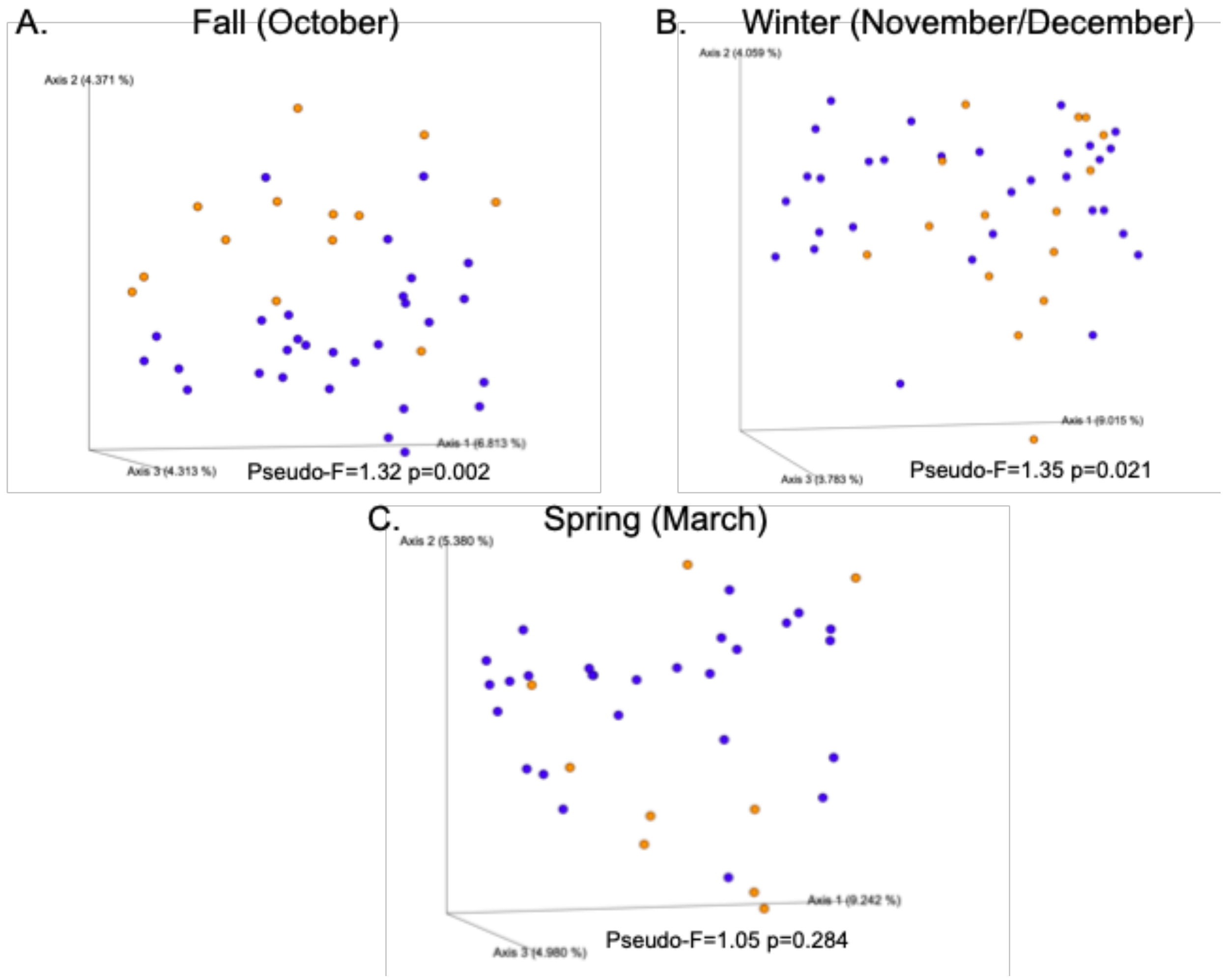

### Additional Figure 3.

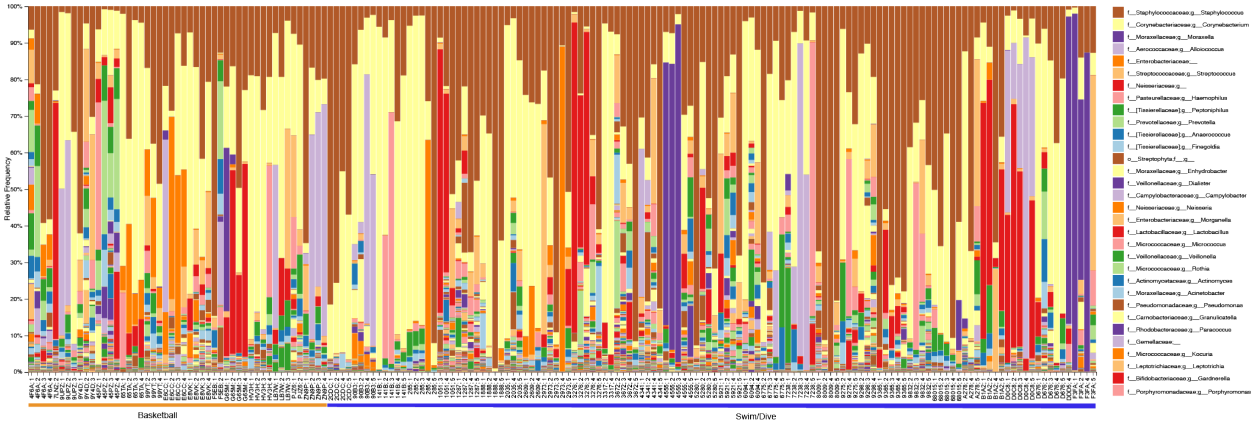


### Additional Figure 4.

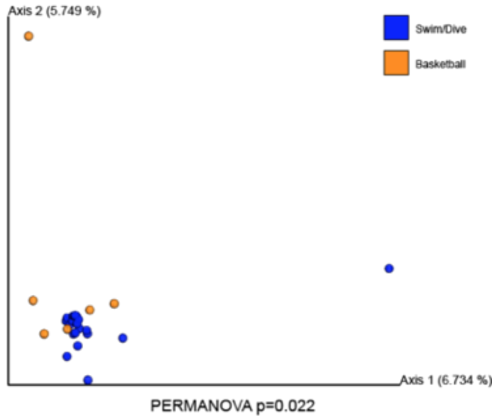

**Additional Figure 5.**

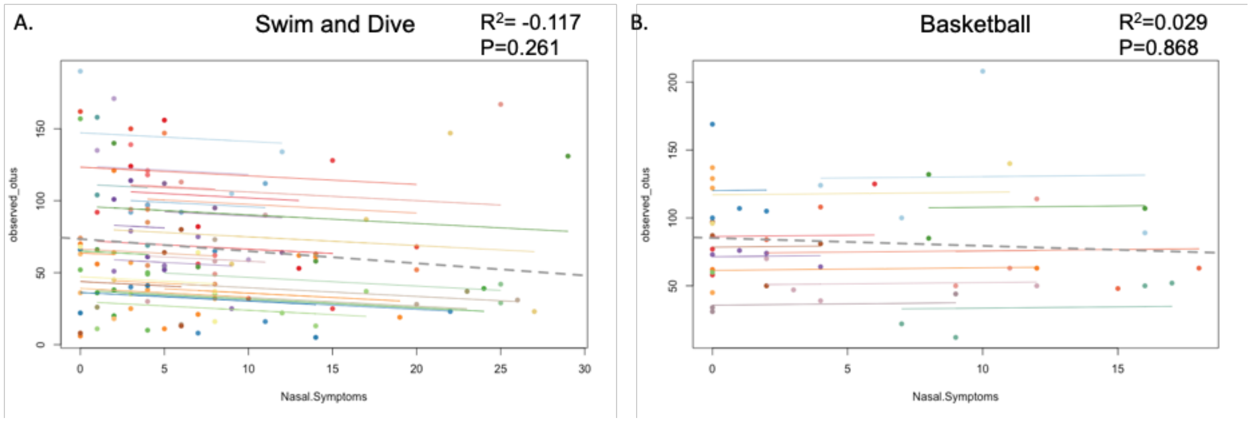


### Additional Figure Legends.

**Additional Figure 1.** Pairwise longitudinal comparisons of observed OTUs in individuals on the Swim/Dive and Basketball teams at each time point. A) Observed OTUs did not change over time in either team. B) Pairwise magnitude of change between sampling timepoints in Basketball and Swim/Dive participants demonstrated no significant changes between any time point pairs tested (p>0.05, Wilcoxon Signed Rank Test).

**Additional Figure 2**. Principal Coordinates analysis of Jaccard dissimilarity index at each timepoint, colored by participants on the Basketball (orange) or Swim/Dive team (blue). In Fall (A) and Winter (B) , significant clustering by team was observed (p<0.05, PERMANOVA). At the last sampling time point (Spring, C), no significant differences were observed between Swim/Dive and Basketball (p=0.284, PERMANOVA).

**Additional Figure 3**. Taxa barplot showing relative abundance of taxa in each individual, sorted by team and individual. The most abundant taxa are *Staphylococcus* (brown) and *Corynebacterium* (yellow). Some individuals have nasal communities dominated by *Moraxella* (purple), *Neisseriaceae* (red), or *Streptococcaceae* (orange).

**Additional Figure 4**. Principal Coordinates Analysis of Non Parametric Microbial Interdependence Network (NMIT) results showing similarities among subjects over time. Each sphere represents an individual’s longitudinal interdependence network over the four sampling time points. These results demonstrate significant differences in longitudinal interdependence networks in each team (p=0.022, pseudo-F=1.23, PERMANOVA).

**Additional Figure 5**. Repeated measures correlation of nasal symptoms and microbiome richness (observed OTUs) in Swim/Dive (A) and Basketball (B) teams estimated using analysis of covariance as implemented in the rmcorr package in R. We did not observe statistical significance for either team (p>0.05).
